# CREPAS: a reproducible nascent chromatin sequencing analysis pipeline for epigenome replication studies

**DOI:** 10.64898/2026.06.21.732899

**Authors:** Samuel Ruiz-Pérez, Qian Du, Alva Biran, Anja Groth, Nicolas Alcaraz

## Abstract

Chromatin-based genomics data are essential for understanding genome regulation and the mechanisms underlying epigenetic memory. Recent methods such as ChOR-seq and SCAR-seq assess histone modifications and chromatin-associated proteins during and after replication, capturing chromatin states that contribute to memory across cell divisions. Current tools for chromatin data analysis lack scalability and reproducibility across computing infrastructures, offer limited parameters, and are applicable only to a few sequencing techniques, ignoring the information from nascent chromatin assays. To address these challenges, we developed CREPAS, a Nextflow pipeline for analyzing nascent and parental chromatin sequencing data, including ChIP-seq, ChOR-seq, SCAR-seq, OK-seq, ATAC-seq, CUT&RUN, and CUT&Tag, and derivative protocols. CREPAS provides an end-to-end solution, from quality control to advanced analyses, including downsampling, peak calling, annotation, and visualization. By harnessing quantitative assays such as qChIP-seq and qChOR-seq, the normalization methods in CREPAS allow to compare the restoration kinetics of individual marks or proteins across replication timepoints. Moreover, the pipeline includes calculations such as fork directionality and partitioning using OK-seq and SCAR-seq data, linking replication dynamics to epigenetic inheritance. CREPAS is a valuable resource that enhances the efficiency and reproducibility of nascent chromatin sequencing data analyses, enabling the study of chromatin replication and propagation of epigenetic states.

**GRAPHICAL ABSTRACT:** 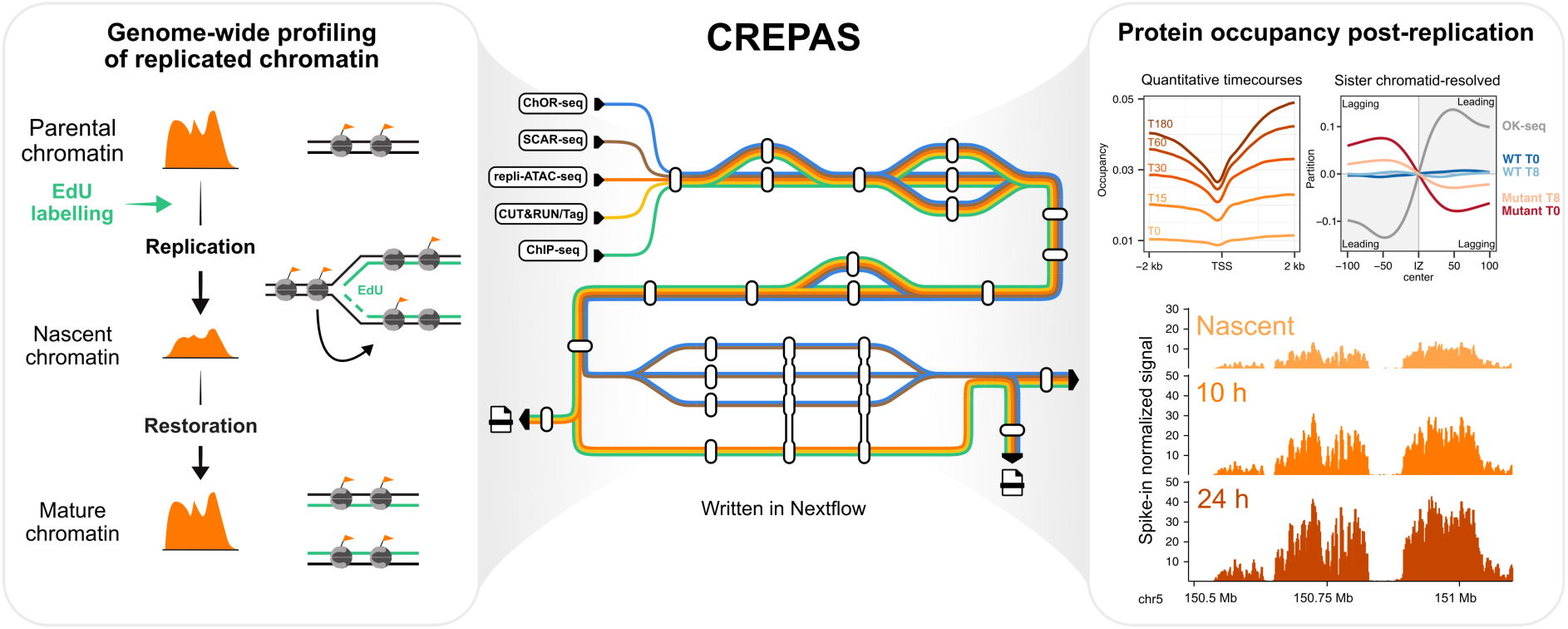

## INTRODUCTION

Epigenetic memory allows cells to preserve stable gene expression programs and respond appropriately to developmental and environmental cues. The establishment and maintenance of this memory rely on chemical modifications to DNA and nucleosomes, which regulate genome accessibility, promote accurate DNA repair, repress transposable elements, and fine-tune transcriptional outputs [1]. Such chromatin features are essential for genome stability and cellular identity, making their faithful maintenance during cell division critical for preventing dysfunction and disease [2,3].

Chromatin presents a physical barrier to the replication machinery and is therefore transiently disrupted during DNA replication. Nucleosomes are disassembled ahead of advancing replication forks and rapidly reassembled on the newly replicated daughter strands [4,5]. Understanding how epigenetic information is transmitted during DNA replication requires methods that capture dynamic and locus-specific chromatin events. Traditional chromatin immunoprecipitation (ChIP) techniques, together with genomic and proteomic readouts, have provided important insight into chromatin organization in steady-state cell populations. However, these approaches do not resolve how chromatin composition is disrupted, transmitted, and restored as replication forks progress [6]. Proteomic methods such as isolation of proteins on nascent DNA (iPOND) [7] and nascent chromatin capture (NCC) [8] have broadened our understanding by characterizing proteins associated with nascent and mature chromatin and by defining global patterns of histone modification dynamics after replication [9]. Despite their utility, these strategies lack the locus-specific resolution required to uncover region-specific mechanisms of chromatin inheritance.

A wide range of sequencing-based methods have further advanced the ability to map replication-coupled chromatin dynamics with locus-level precision [10]. Through labeling of nascent DNA with 5-ethynyl-2′-deoxyuridine (EdU) and digestion of chromatin with micrococcal nuclease, nascent chromatin avidin pull-down (NChAP) has revealed that nucleosome positioning is rapidly established after DNA replication [11]. Replication-coupled assay for transposase-accessible chromatin (repli-ATAC-seq) has been used to monitor the restoration of chromatin accessibility, revealing that transcription restart plays a key role in re-establishing open chromatin states on nascent DNA [12].

Similarly, chromatin occupancy after replication (ChOR-seq) has enabled genome-wide profiling of protein occupancy on replicated DNA, providing insight into how chromatin-associated factors are restored on nascent DNA [6]. ChOR-seq entails a ChIP followed by isolation of EdU labeled DNA fragments by a biotin Click-iT reaction and streptavidin pulldown (**Figure 1A**) [6]. In a ChIP-seq experiment, the shearing, amplification, and sequencing steps can introduce technical and lead to artifactual signal enrichment across the genome. Thus, not all sequenced DNA fragments represent true protein-binding events. To control against this background signal and to distinguish true protein-binding peaks from noise during downstream analyses, a sample collected after shearing but before the immunoprecipitation step is commonly used (*chromatin input control*, also called *total input*). Similarly, in ChOR-seq, the EdU-labeled fraction of the chromatin input is used as an additional control to account for both the efficiency and genome coverage of EdU labeling (*EdU input control*, also referred to as *click input*).

**Figure 1.**
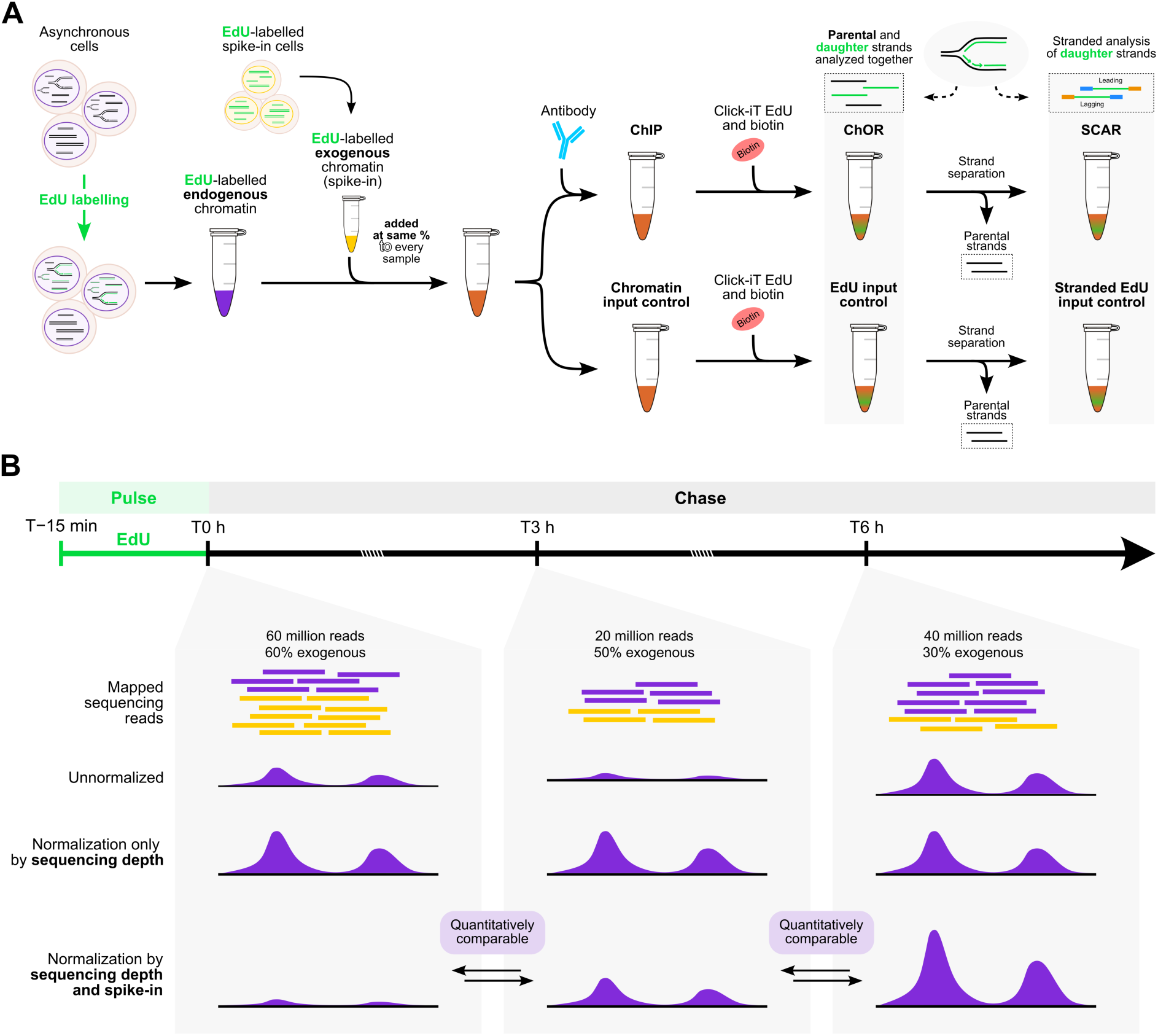
Overview of experimental steps in ChOR-seq and SCAR-seq and spike-in normalization methods for quantitative comparisons of occupancy across DNA replication timepoints. **(A)** Both ChOR-seq and SCAR-seq methods are based on a common workflow that combines EdU pulse labeling of newly synthesized DNA with chromatin immunoprecipitation, followed by purification and high-throughput sequencing. ChOR-seq enables quantitative profiling of histone modifications and chromatin-associated proteins after replication, and SCAR-seq distinguishes chromatin occupancy between the two nascent sister chromatids. **(B)** By adding an equal amount of exogenous spike-in chromatin to each sample in a qChOR-seq time course, the sequential ChIP and EdU-mediated enrichment of replicated DNA produces the same absolute recovery of spike-in DNA across samples. This enables the normalization of endogenous ChIP read counts and direct quantitative comparisons across samples.

ChOR-seq allows to quantitively assess protein occupancy across multiple timepoints after DNA replication, for example, in studies of histone PTM restoration. This is achieved through a pulse-chase EdU-labeling time course where the nascent sample is compared with progressively more mature samples (**Figure 1B**). To enable quantitative comparison across these different time points (qChOR-seq), an equal amount of exogenous EdU-labeled spike-in chromatin (e.g., from *Drosophila melanogaster*) must be added to all samples prior to the ChIP step. After sequencing, read coverage is normalized to the total number of exogenous reads. Because a constant amount of reference chromatin is added to each sample, the sequential ChIP and EdU-mediated pulldown results in the same absolute enrichment of the spike-in DNA across samples, whereas the absolute enrichment of endogenous DNA remains sample-specific. This normalization therefore enables direct comparison between samples and helps correct for the experimental and analytical limitations of traditional ChIP-seq.

Overall, sequencing-based assays such as NChAP, repli-ATAC-seq, qChIP-seq and qChOR-seq have helped define how nucleosome organization, chromatin accessibility, transcription factor occupancy, and protein binding are re-established genome-wide following replication fork passage.

Another major advance came with the development of strand-specific sequencing-based methods that distinguish chromatin assembled on the leading and lagging strands of replication. Techniques such as NChAP [11], sister chromatids after replication (SCAR-seq) [6], and enrichment and sequencing of protein-associated nascent DNA (eSPAN) [13] isolate nascent and parental DNA strands, enabling quantitative analysis of strand-biased occupancy of histones or chromatin-associated factors. In SCAR-seq, replicated DNA labelled with EdU is biotinylated and captured using streptavidin beads. The DNA is then denatured to remove the unlabeled parental strands, leaving only the newly synthesized EdU-labeled strands (**Figure 1A**). These nascent strands are subsequently amplified and sequenced, thereby preserving strand-specific information [6]. Genome-wide associations with the leading and lagging strand can be inferred from replication origins in yeast [14,15] or from replication fork direction maps, replication initiation zones or origins defined by Okazaki fragment sequencing (OK-seq) [16,17], EdU and hydroxyurea and sequencing (EdUseq-HU) [18], or single-cell EdU sequencing (scEdU-seq) [19] in mammalian cells. These approaches have been instrumental in identifying the contributions of replisome components and chromatin regulators to replication-coupled epigenome maintenance and for characterizing genome-wide chromatin maturation at strand resolution. Together, these sequencing-based methods provide a rich foundation for understanding how chromatin features are disrupted and faithfully restored during DNA replication, and they motivate the development of computational tools capable of analyzing these diverse datasets.

An unprecedented volume of data across cell types, experimental modalities, and biological contexts has been generated with chromatin sequencing assays to study the regulation of the genome and epigenome. Despite the informational richness of these datasets, their full potential remains underexploited, as fragmentation at the analysis stage results in chromatin data being processed through heterogeneous, assay-specific pipelines that vary in preprocessing, normalization, quality control, and parameterization, thereby limiting comparability across studies and laboratories. Widely used chromatin segmentation methods such as ChromHMM [20] and Segway [21] have demonstrated that chromatin features are highly informative for distinguishing functional genomic states, but they rely on consistently processed and uniformly parameterized input data, including fixed binning and standardized signal definitions, assumptions that are often violated across independently processed datasets.

In practice, current tools for chromatin data analysis generally lack the scalability and reproducibility required for execution across diverse high-performance computing environments. Moreover, most available pipelines are distributed as *ad hoc* collections of scripts rather than as modular, well-structured, and containerized software, which hinders transparent versioning, dependency control, and reproducible deployment. As a result, replicating published analyses is often challenging. Existing pipelines also present limited parameters and are not designed to accommodate the distinctive characteristics of nascent chromatin sequencing datasets (**Table 1**), restricting their ability to fully capture histone-mark dynamics, protein occupancy, and chromatin accessibility across DNA replication.

**Table 1.** Comparison of available pipelines for DNA sequencing analysis.

| Pipeline | Workflow language | Features |  |  |  |  |  |  |  |  |
| --- | --- | --- | --- | --- | --- | --- | --- | --- | --- | --- |
|  |  | Multi-assay/omics | Peak calling | Bin/window-based quantification | UMI-based deduplication | Downsampling | Spike-in normalization | Repetitive element read allocation and quantification | ENCODE ChIP-seq/ATAC-seq standards | Replication fork directionality and partition analyses |
| CREPAS | Nextflow | ✓ | ✓ | ✓ | ✓ | ✓ | ✓ | ✓ | ✓ | ✓ |
| nf-core/chipseq 2.1.0 [22] | Nextflow |  | ✓ | ✓ |  |  |  |  |  |  |
| nf-core/atacseq 2.1.2 [23] | Nextflow |  | ✓ | ✓ |  |  |  |  |  |  |
| nf-core/cutandrun 3.2.2 [24] | Nextflow |  | ✓ |  |  |  | ✓ |  |  |  |
| GenPipes/ChIP-Seq 6.1.0 [25] | Python |  | ✓ | ✓ |  |  |  |  |  |  |
| ENCODE 3 ChIP-seq pipeline 2.2.2 [26] | Caper/Cromwell |  | ✓ |  |  | ✓ |  |  | ✓ |  |
| SnakePipes 3.3.0 [27] | Snakemake | ✓ | ✓ | ✓ | ✓ |  | ✓ |  |  |  |
| ePeak 1.0 [28] | Snakemake |  | ✓ |  |  |  | ✓ |  |  |  |

Here, we present CREPAS, a Nextflow DSL2 pipeline for processing and analyzing nascent (ChOR-seq, SCAR-seq, eSPAN, repli-ATAC-seq, Okazaki-seq) and parental (ChIP-seq, CUT&RUN, CUT&Tag, ChIP-exo, ATAC-seq, DNAse-seq) epigenomic sequencing data. We demonstrate the efficiency and reproducibility of this pipeline through two applications using ChOR-seq and SCAR-seq datasets spanning various histone marks. Rather than functioning as another assay-specific workflow, this pipeline is designed as a comprehensive and extensible framework with the potential to become a standard solution for chromatin sequencing data analysis. Altogether, we highlight how CREPAS enables robust and quantitative analyses of nascent DNA sequencing data to study replication-coupled chromatin dynamics and epigenome maintenance.

## MATERIALS AND METHODS

### Implementation

CREPAS is written in Nextflow [29] and has been tested with versions > 24.10.0. By leveraging Nextflow’s domain-specific language version 2 (DSL2), the pipeline benefits from a fully modular architecture with clearer organization and improved maintainability. Moreover, the pipeline is built on the nf-core template, which provides standardized coding framework and comprehensive documentation that ensure current best practices [30,31]. Most modules used in CREPAS are available in the nf-core/modules [31] community repository, which provides Nextflow wrappers around individual tools. These tools are distributed via (bio)conda [32] packages and are accompanied by a corresponding Docker [33] and Singularity [34] container provided by the Biocontainers [35] community or the Seqera Containers (https://seqera.io/containers/) infrastructure, enabling portability and reproducibility for each module. This single-tool-per-process approach ensures that dependency conflicts are mitigated [36]. Thanks to CREPAS’s structure, which mostly adheres to the nf-core guidelines [31], contributors can also use helper packages such as nf-core/tools [31] to easily create, install and re-use new modules to extend the functionalities of CREPAS, similar to what has been achieved with other pipelines [36].

### Sequencing datasets

To test the pipeline’s performance on high-resolution nascent chromatin datasets, we selected published quantitative ChOR-seq (qChOR-seq) data of H3K27me3 and H2BK120ub1 [37]. We then extended this evaluation to strand-specific data by applying CREPAS to published SCAR-seq datasets for H3K27me3 and H3K27ac [38], thereby enabling the assessment of complementary nascent chromatin profiling modalities. The datasets analyzed in this study are summarized in **Table 2**.

**Table 2.**
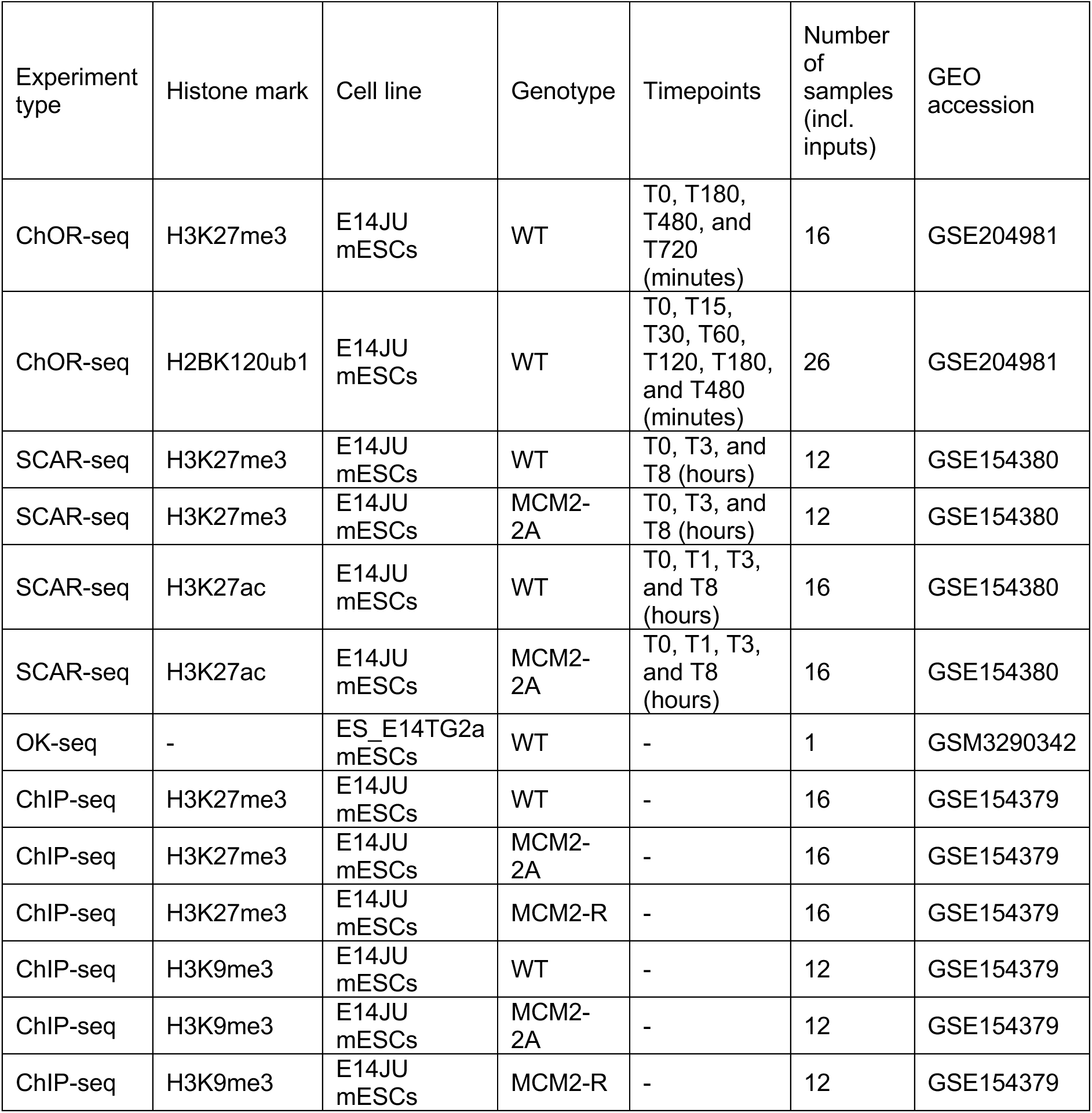
Summary of nascent chromatin sequencing datasets analyzed with CREPAS in this study.

### Compute environment and pipeline execution

All pipeline runs were executed on the DAN System high-performance computing (HPC) cluster at the Novo Nordisk Foundation Center for Stem Cell Medicine (reNEW), Faculty of Health and Medical Sciences, University of Copenhagen. Each of the experiments (one per assay type and histone mark) was run independently, with a primary Nextflow job initiated on the head (submission) node using 2 CPUs and 4 GB of memory. The head node is equipped with 6 virtual CPUs and 12 GB of RAM and operates on Red Hat Enterprise Linux 8.10 with the SLURM 20.11.9 workload manager. All downstream pipeline processes were executed on a dedicated compute node configured with two AMD EPYC 7763 processors (256 hyper-threaded cores) and 4 TB of RAM, also running Red Hat Enterprise Linux 8. These jobs were submitted using the cluster-specific configuration file (ku_sund_danhead.config), which limited execution to a maximum of five concurrent jobs, each permitted up to 8 CPUs, 64 GB of RAM, and a 72-hour maximum runtime. The workflow was executed using Nextflow 25.10.2 [29], running under OpenJDK 20.0.0 with Singularity 3.8.7 as the containerization engine.

## RESULTS

### Pipeline feature overview

An overview of the CREPAS pipeline is shown in **Figure 2**. The input data is a standardized samplesheet in comma-separated value (CSV) format, that provides all relevant metadata needed for the analysis as well as the paths to the FASTQ files. The samplesheet and supplied parameters are validated with the nf-schema plugin [39]. Through this samplesheet, the pipeline can handle single-end and paired-end samples, as well as samples for which a separate FASTQ file containing unique molecular identifier (UMI) barcodes is provided. The pipeline can process biological and technical replicates for both samples and input controls of several sequencing techniques, including ChIP-seq [40], ChOR-seq [6], ChIP-exo [41], SCAR-seq [6], OK-seq [16], ATAC-seq [42], CUT&RUN [43], CUT&Tag [44], and TIP-seq [45]. It consists of twelve main processing units:

**Figure 2.**
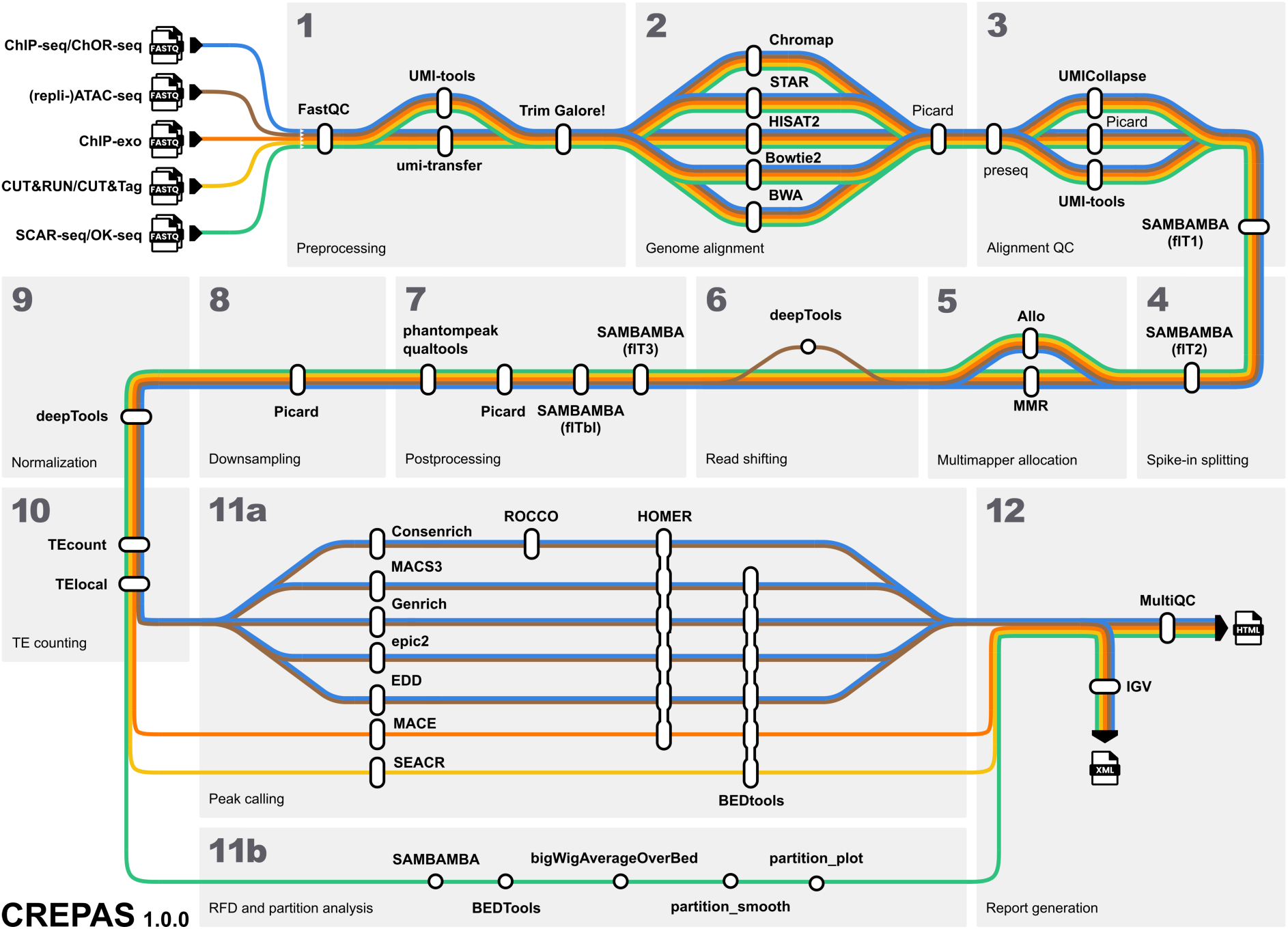
Overview of CREPAS. The pipeline is composed of several subworkflows and modules that handle sequencing read preprocessing, genome alignment, alignment quality control, multimapper allocation, spike-in normalization, downsampling, and transposable element quantification. It also includes downstream analyses tailored to specific applications, such as peak calling, replication fork directionality analysis, and partitioning analyses. Throughout the workflow, the pipeline generates a range of diagnostic plots and summary tables, which are compiled into a MultiQC report and genomic tracks for further data exploration. In addition, several steps offer users a choice between different processing tools (summarized here), enabling efficient benchmarking of both software and parameter settings.

### Pre-processing

The initial raw read quality control steps are performed at the library (technical replicate) level, e.g., when the same library has been sequenced multiple times to increase sequencing depth. First, FastQC [46] is used to generate general quality metrics, including quality score distributions, sequence content, adapter contamination and overrepresented sequences.

Next, CREPAS can handle UMI barcodes, by using either UMI-tools [47] to extract them from the read FASTQ and add them to their corresponding read names, or umi-transfer [48] to transfer the UMIs from a separate FASTQ file to the read FASTQs. Additionally, the user can opt out of extraction/transfer in cases where UMIs have been processed beforehand or are not available.

The raw reads are then passed to Trim Galore! [49], which is a wrapper tool that integrates Cutadapt [50] and FastQC [46] to perform consistent adapter and quality trimming of FASTQ files. By default, it automatically detects and removes the appropriate adapter sequences. However, through parameters in CREPAS, Trim Galore! can be configured to search for the full length (33 bp) TruSeq (ChIP-seq, ChOR-seq, SCAR-seq) or Nextera (ATAC-seq) adapter sequences for both complement and reverse complements. In addition, we have implemented an extra hard-trimming step, in which reads are shortened to a user-specified length from their 3’ or 5’ end. Importantly, this is performed after adapter/quality trimming and can be used to benchmark tools that are affected by a shorter input read length, for example for multimapping read allocation algorithms [51].

### Read mapping

Adapter-trimmed reads are then mapped to a reference genome using the aligner defined by parameter *--aligner*, with available options including BWA-MEM [52], BWA-MEM2 [53], Chromap [54], Bowtie [55], Bowtie2 [56], STAR [57], HISAT2 [56], Minimap2 [58] or strobealign [59]. Each aligner requires a genome index, which can either be provided via the corresponding parameter (e.g., *--chromap_index*) or generated automatically from the input genome FASTA file. Since index generation can be time-intensive for large genomes, the *--save_reference* option allows users to store the created indices for future runs, thereby reducing processing time.

The pipeline is designed so that all files generated after alignment are stored in a directory corresponding to the aligner specified with the *--aligner* parameter. For example, if *--aligner chromap* is used, all subsequent results will be saved in the *chromap/* directory. This structure keeps outputs organized and enables users to easily obtain results from multiple aligners by re-running the pipeline with a different *--aligner* option alongside the *-resume* flag, without overwriting existing data. This setup also facilitates benchmarking across alignment algorithms. Similarly, we have implemented this output directory structure for all the downstream analyses of the pipeline, so that the user can test several tools for each step without compromising the output files of previous runs.

### Post-alignment quality control

After mapping, if more than one technical replicate has been provided for a specific sample (biological replicate), then these library-level alignments are merged with Picard MergeSamFiles [60] and subsequently used for the downstream analyses.

Next, duplicate reads can be removed from the aligned datasets to account for library fragments that may have been sequenced multiple times because of PCR amplification biases. The pipeline can perform UMI-based deduplication using UMI-tools dedup [47], or UMICollapse [61], which implements many efficient algorithms for faster UMI deduplication. Otherwise, if UMIs have not been provided to the pipeline, duplicates can be marked in the alignment files using Picard MarkDuplicates [60] and filtered in the following processing steps.

During the first filtering step, Sambamba view [62] is used to remove unmapped or improperly paired reads and reads marked as duplicates. Then, if reads were mapped to a hybrid reference genome (e.g., mm39 with dm6) to account for spike-in content in the samples, the alignments are split by genome and filtered again using Sambamba [62]. Thus, this step generates one alignment file containing endogenous reads and another containing exogenous (spike-in) reads.

Multimapping reads (MMRs) have historically been a challenge in genomic analyses, particularly in ChIP-seq experiments, where their exclusion can lead to incomplete or biased results [63]. Various methods have been developed to address the challenges of MMRs in sequencing data. Some allocate MMRs by leveraging uniquely mapped read counts or coverage densities near potential mapping sites [64], while more recent approaches perform probabilistic allocation of MMRs using convolutional neural network models [51]. With CREPAS, multimapper allocation can be carried out on both endogenous and exogenous reads with MMR [65] or Allo [51], which has been trained and tested on a wide range of transcription factor and histone modification data. Alternatively, MMRs can be allocated as early as the mapping step with the multimapping read allocation method implemented in Chromap [54].

Afterwards, a third filtering step using Sambamba [62] is performed on the processed alignments to remove unmapped or improperly paired reads, secondary alignments, and reads with low mapping quality (depending on the aligner used). Additionally, the user of CREPAS can provide a file with blacklisted regions to remove reads that were mapped to locations that usually generate artifactual, or abnormally high signals in high-throughput sequencing experiments [66].

### Downsampling and quantitative normalization

Next, Picard DownsampleSam [60] can be used to apply a downsampling algorithm to the processed BAM files to retain only a (deterministically random) subset of the reads. Reads from the same template (e.g. read-pairs, secondary and supplementary reads) are all either kept or discarded as a unit. The probability of a read being contained in the subsampled file will depend on the selected downsampling method, which can be chosen to account for endogenous or exogenous (spike-in) reads across samples or input controls.

In addition, in CREPAS we have implemented various parameters to modify the conditions on which the downsampling probabilities are calculated, including thresholds of minimum endogenous or exogenous reads. For example, in the case of downsampling methods that search for the minimum endogenous mapped reads across a set of samples, the *--downsampling_endo_threshold* parameter (e.g., 10000000 reads) will determine the lower limit of that minimum. Similarly, for downsampling methods that search for the minimum exogenous mapped reads across a set of samples, the *--downsampling_exo_threshold* parameter (e.g., 300000 reads) will determine the lower limit of that minimum. If there are no totals above this threshold, then the (true) minimum is used. We have also added a parameter that allows the user to provide a list of histone marks for which the downsampling probabilities should be calculated using the total number of reads before the low-quality filtering performed in the previous processing steps. For example, for ChIP-seq samples of H3K9me3 or H3K27me3, the user might prefer to downsample considering the total mapped reads including multimapping reads, which are otherwise unaccounted for at this step.

Coverage tracks can then be generated for the final filtered, and optionally, downsampled, BAM files with deepTools bamCoverage [67]. The coverage is calculated as the number of reads per bin (*α_i_*), where bins are short consecutive counting windows of a defined size. CREPAS provides various ways of using quantitative chromatin sequencing methods by normalizing the calculated coverage to account for input control or spike-in reads. These normalization methods are described in **Table 3**. In the case of quantitative ChOR-seq experiments, an equal amount of exogenous spike-in EdU-labeled chromatin is added to all samples before the ChIP step to allow for the quantitative comparison between different time points [68]. After sequencing, results can be normalized to the total number of exogenous reads (here denoted as SRPM) [68], allowing the quantitative assessment of the restoration kinetics of individual marks or proteins to pre-replication levels [6]. An alternative strategy for spike-in normalization additionally corrects for minor variations in added spike-in chromatin in different samples by using the ratio of exogenous to endogenous total read counts in the corresponding input samples (here denoted as CISRPM) [69]. Before CISRPM normalization, an extra downsampling step, either with respect to the endogenous or exogenous reads, may be applied as another way to even out minor variations between samples. Moreover, to assess the relative signal over background for each location in the genome, the ratio of the IP CISRPM and input CISRPM values can be calculated per genomic bin (CISRPM SOI).

**Table 3.** Summary of normalization methods implemented in CREPAS that account for endogenous and exogenous reads in both ChIP/ChOR and input controls.

| Method | Description | Formula | References |
| --- | --- | --- | --- |
| Raw | No normalization | $\alpha_{\text{Raw}} = \alpha_i \times 1$ | - |
| RPM | Reads Per Million mapped reads | $\alpha_{\text{RPM}} = \alpha_i \times \frac{10^6}{\text{total mapped reads}}$ | - |
| SRPM | Spike-in-normalized Reads Per Million mapped reads | For the endogenous ChIP:<br>$\alpha_{\text{SRPM}_i} = \alpha_i \times \frac{10^6}{\frac{\text{total mapped}}{\text{exogenous ChIP reads}}}$<br>For the endogenous chromatin input control:<br>$\alpha_{\text{SRPM}_i} = \alpha_i \times \frac{10^6}{\frac{\text{total mapped}}{\text{exogenous chromatin input control reads}}}$<br>For the endogenous ChOR:<br>$\alpha_{\text{SRPM}_i} = \alpha_i \times \frac{10^6}{\frac{\text{total mapped}}{\text{exogenous ChOR reads}}}$<br>For the endogenous EdU input control:<br>$\alpha_{\text{SRPM}_i} = \alpha_i \times \frac{10^6}{\frac{\text{total mapped}}{\text{exogenous EdU input control reads}}}$ | [6,68] |
| CISRPM | ChIP/ChOR-and-Input-Spike-in-normalized Reads Per Million mapped reads | For the endogenous ChIP:<br>$\alpha_{\text{CISRPM}_i} = \alpha_i \times \frac{10^6}{\frac{\text{total mapped}}{\text{exogenous ChIP reads}}} \times \frac{\frac{\text{total mapped}}{\text{exogenous chromatin input control reads}}}{\frac{\text{total mapped}}{\text{endogenous chromatin input control reads}}}$<br>For the endogenous chromatin input control:<br>$\alpha_{\text{CISRPM}_i} = \alpha_i \times \frac{10^6}{\frac{\text{total mapped}}{\text{exogenous chromatin input control reads}}} \times \frac{\frac{\text{total mapped}}{\text{exogenous chromatin input control reads}}}{\frac{\text{total mapped}}{\text{endogenous chromatin input control reads}}} = \alpha_{\text{RPM}}$<br>For the endogenous ChOR:<br>$\alpha_{\text{CISRPM}_i} = \alpha_i \times \frac{10^6}{\frac{\text{total mapped}}{\text{exogenous ChOR reads}}} \times \frac{\frac{\text{total mapped}}{\text{exogenous EdU input control reads}}}{\frac{\text{total mapped}}{\text{endogenous EdU input control reads}}}$<br>For the endogenous EdU input control:<br>$\alpha_{\text{CISRPM}_i} = \alpha_i \times \frac{10^6}{\frac{\text{total mapped}}{\text{exogenous EdU input control reads}}} \times \frac{\frac{\text{total mapped}}{\text{exogenous EdU input control reads}}}{\frac{\text{total mapped}}{\text{endogenous EdU input control reads}}} = \alpha_{\text{RPM}}$ | [37,69] |
| CISRPM-SOI | CISRPM Signal Over Input | For ChIP:<br>If<br>$\alpha_{\text{CISRPM}_{\text{ChIP}_i}} > \text{--min\_signal\_for\_soi},$<br>and<br>$\alpha_{\text{CISRPM}_{\text{Chromatin input}_i}} > \text{--min\_input\_for\_soi},$<br>then:<br>$\alpha_{\text{CISRPM-SOI}_i} = \frac{\alpha_{\text{CISRPM}_{\text{ChIP}_i}}}{\alpha_{\text{CISRPM}_{\text{Chromatin input}_i}}}$<br>Otherwise:<br>$\alpha_{\text{CISRPM-SOI}_i} = \text{NaN}$<br>For ChOR:<br>If<br>$\alpha_{\text{CISRPM}_{\text{ChOR}_i}} > \text{--min\_signal\_for\_soi},$<br>and<br>$\alpha_{\text{CISRPM}_{\text{EdU input}_i}} > \text{--min\_input\_for\_soi},$<br>then:<br>$\alpha_{\text{CISRPM-SOI}_i} = \frac{\alpha_{\text{CISRPM}_{\text{ChOR}_i}}}{\alpha_{\text{CISRPM}_{\text{EdU input}_i}}}$<br>Otherwise:<br>$\alpha_{\text{CISRPM-SOI}_i} = \text{NaN}$ | This study |

### Counting transposable element reads

Transposable elements (TEs) are mobile DNA sequences that make up a substantial portion of most eukaryotic genomes. In mammals, ChIP-seq studies have revealed that for any given transcription factor and cell type examined, TEs contribute a substantial fraction of binding sites across the genome (5–40%; average ∼20%) [70–72]. Despite their abundance, reads associated with TEs are frequently excluded from sequencing data analyses due to the difficulty of accurately assigning ambiguously mapped reads to specific loci. Various strategies have been proposed to mitigate this problem, such as discarding MMRs altogether [73,74]. While this approach enhances analytical specificity, it also results in the loss of a significant fraction of TE-derived reads. Therefore, tools such as TEtranscripts/TEcount [74] and TElocal [75] have been developed to analyze both gene- and TE-associated reads concurrently, specifically in gene/transcript expression analysis, but can also be used on chromatin sequencing data. In the CREPAS pipeline, the user has the option to run both TEcount and TElocal on the processed alignment files.

We tested TE quantification on ChIP-seq data of H3K9me3 and H3K27me3, from a study that mutated the histone-binding domain of MCM2, a component of the replicative helicase [38]. This mutation prevents the protein from binding parental H3–H4 and disrupts the faithful transmission of histone marks, leading to a loss of fidelity in the H3K9me3 landscape and a global gain of H3K27me3 [38]. Mutant (MCM2-2A) cells show a redistribution of H3K9me3 signal, with a major loss from repetitive regions and a gain in the nonrepetitive genome. Consistent with this, we found a decreased fraction (∼28% to ∼22%) of multimapping reads in the H3K9me3 datasets, and a minor increase in multimapper fraction in H3K27me3 (**Supplementary Figures S1A and S1B**). Importantly, the inclusion of multimapping reads in quantification and differential occupancy analysis led to an increase in the number of detected TE families with loss of H3K9me3 (**Supplementary Figure S1C**). In contrast, including multimappers reduced the number of detected TE families with an increase in H3K27me3, possibly due to an improvement in false discovery rate control (**Supplementary Figure S1D**).

### Peak calling

CREPAS includes a comprehensive set of peak callers to extract domains of enrichment with different properties depending on the organism, condition, biological feature of interest, or sequencing protocol. The 9 included tools are MACS3 [76], Genrich [77], epic2 [78], ROCCO [79], DANPOS2 [80], MACE [81], SEACR [82], phantompeakqualtools SPP [83], and EDD [84]. Additionally, Consenrich [85] can be used upstream of ROCCO to generate genome-wide, uncertainty-calibrated signal tracks from noisy multi-sample data and detect a consensus of open chromatin regions, TF footprints, nucleosome occupancy, or histone modifications. Other tools could be added with relatively low effort thanks to the modularized structure of the pipeline. While some peak callers are recommended for specific protocols, such as MACE for ChIP-exo, the user can select any set of tools to be run parallelly on their data. This facilitates the benchmarking of peak calling algorithms, while also ensuring a low resource footprint by only running the desired tools.

The pipeline also is able to generate and run irreproducible discovery rate (IDR) [86] analysis on all pairs of replicates, self-pseudoreplicates and pooled pseudoreplicates in a parallelized manner. This allows to generate consensus and reproducible peak sets, together with the computation of several quality metrics for each sample or group of samples, including rescue and self-consistency ratios, fraction of reads in peaks (FRiP) and cross-correlation scores. Moreover, it allows the user to evaluate compliance to the working standards and guidelines for ChIP experiments of the ENCODE and modENCODE consortia [83].

### Fork directionality and partition signal analyses

Based on the previously published protocols for OK-seq and SCAR-seq, CREPAS can handle OK-seq data to derive replication fork directionality (RFD) [16,17], and SCAR-seq data to yield histone partition signals [6]. Both RFD and partition signal computation starts with the mapped and processed alignments, which are then split into reads mapping to the forward and reverse strands. Each strand’s read files are converted to bigWig coverage tracks using BEDTools genomecov [87], and reads mapping to either strand are counted in book-ended bins of 1 kb across the genome using bigWigAverageOverBed [88]. Importantly, in this step bin and step sizes can be modified through parameters in CREPAS to generate disjoint or sliding bins depending on the needs of the user. Next, each strand’s average counts are normalized to the total mapped reads per million (RPM) with a pseudocount of 1.

In CREPAS, the RFD is calculated from OK-seq reveals the proportions of forks moving rightward (positive values, replicating forward strand by leading strand mechanism) and leftward (negative values, replicating reverse strand by leading strand mechanism) within each window (**Figure 3A**) [17]. The partition value from SCAR-seq indicates the ratio of histones with a specific modification being segregated to the newly synthesized forward (positive values) and newly synthesized reverse (negative values) strand within each window (**Figure 3B**), i.e., the relative difference in histone modification abundance between the two sister chromatids in a cell population [6,17]. Next, partition signal from SCAR-seq and RFD signal from OK-seq are smoothed similarly for visualization purposes. Smoothing is computed across the book-ended 1 kb bins generated in the previous bigWigAverageOverBed step, by performing a uniform blur considering the neighboring 30 bins on each side. In CREPAS, the default smoothing radius of 30 bins can also be adjusted by the user according to the noise level and desired visual resolution. For example, it can be increased to 80 bins for genome views of H4K5ac, H4K20me2, and H3K36me3 SCAR-seq samples [17].

**Figure 3.**
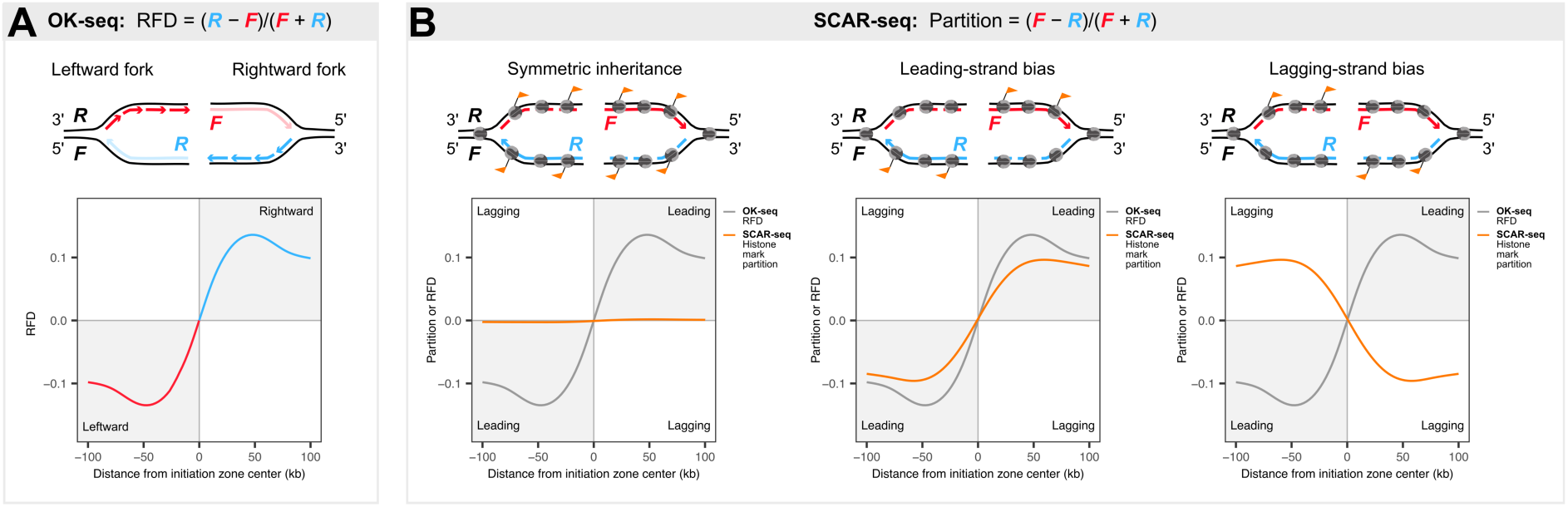
Principle of OK-seq and SCAR-seq for replication fork directionality (RFD) and partition analyses. **(A)** Schematic of leftward and rightward forks with detected with Okazaki fragment sequencing and expected summary RFD profile around initiation zones. (**B**) Schematics and SCAR-seq partition summary plots of histone mark with (left) symmetric inheritance, (center) inheritance biased to the leading strand, and (right) inheritance biased to the lagging strand. Grey curve corresponds to RFD from OK-seq and orange curve represents histone mark partition from SCAR-seq.

Subsequently, OK-seq data can be used to detect initiation zones. CREPAS implements a nonparametric derivative estimation of RFD signal on the genomic bins to detect sharp transitions between neighboring bins, defining replication initiation zones (ascending slopes) [17,89]. These initiation zones are then filtered further. Only bins 1) with normalized counts above a threshold (e.g., *RPM* > 3), 2) with RFD derivative above the 0.9 quantile and zero-crossing, and 3) located within autosomes (e.g., chr1-chr19 for mouse) are kept. Additionally, for initiation zones less than 3 bins apart, the bin with the highest RFD derivative is selected. All these cutoff thresholds for initiation zone definition can be modified by the user through parameters in CREPAS. Then, for visualization purposes, RFD rates around the defined initiation zones are computed and plotted up to, e.g., 100 kb upstream and 100 kb downstream of each initiation zone by averaging the RFD values within each of the 100 bin positions.

Conversely, SCAR-seq data can be used for partition skew detection. In this case, the nonparametric derivative estimation is used on the partition signal to detect sharp transitions. For visualization, the partition rates can then be calculated and plotted up to, e.g., 100 kb upstream and 100 kb downstream of each initiation zone by averaging the partition values within each of the 100 bin positions. Plotting partition rates requires that the user provides defined initiation zones, either as OK-seq data or as intervals in a BED file.

### The spike-in normalization in CREPAS enables quantitative comparisons of histone mark occupancy across post-replication timepoints

We present a standard use case of CREPAS on ChOR-seq data of two different histone marks, H3K27me3 and H2BK120ub1, to showcase the utility of spike-in normalization for studying mark restoration patterns post-replication. The H3K27me3 samples encompassed a pulse-chase EdU-labeling time course, with two biological replicates per timepoint (T0, T180, T480, and T720 hours), each with its corresponding EdU input control (see **Methods**). Additionally, the ChOR samples were resequenced to increase the depth, resulting in 18 samples in total. Parallelly, to test if CREPAS is able to reproduce the results of previous work on different histone marks, we analyzed published ChOR-seq data of the active mark H2BK120ub1 [33]. These included 32 samples, with two biological replicates per timepoint (T0, T15, T30, T60, T120, T180, and T480 minutes), each with its corresponding EdU input control.

As reported in previous studies [37,90], we found that H3K27me3 ChOR-seq signal gradually accumulated across the three timepoints, with the major increase taking place between 3 h and 8 h after S phase (**Figure 4A**). Conversely, H2BK120ub1 showed fast kinetics with full restoration in 120–180 min and prior to cell division (**Figure 4B**) [37]. CREPAS can also generate occupancy profile plots over both consensus peaks and genes (**Figure 4A-B**). For H3K27me3, signal around peak centers increased in both width and height, with broader domains 8 h after replication (**Figure 4A**). In the case of H2BK120ub1, consensus peak centers were clearly demarcated and maintained at all time points (**Figure 4B**) [37]. With the output of the pipeline, we further characterized H2BK120ub1 signal at TSSs, which contain an upstream nucleosome-depleted region (NDR) to facilitate transcription [91]. As previously reported, nascent and maturing H2BK120ub1 signals at TSSs were clearly restricted to regions downstream of the TSS, supporting the observation of accurate recycling and restoration (**Figure 4B**) [37]. H3K27me3 profiles also displayed well-defined NDRs that remained stable over time, though their boundaries appeared less sharp in nascent chromatin (**Figure 4A**).

**Figure 4.**
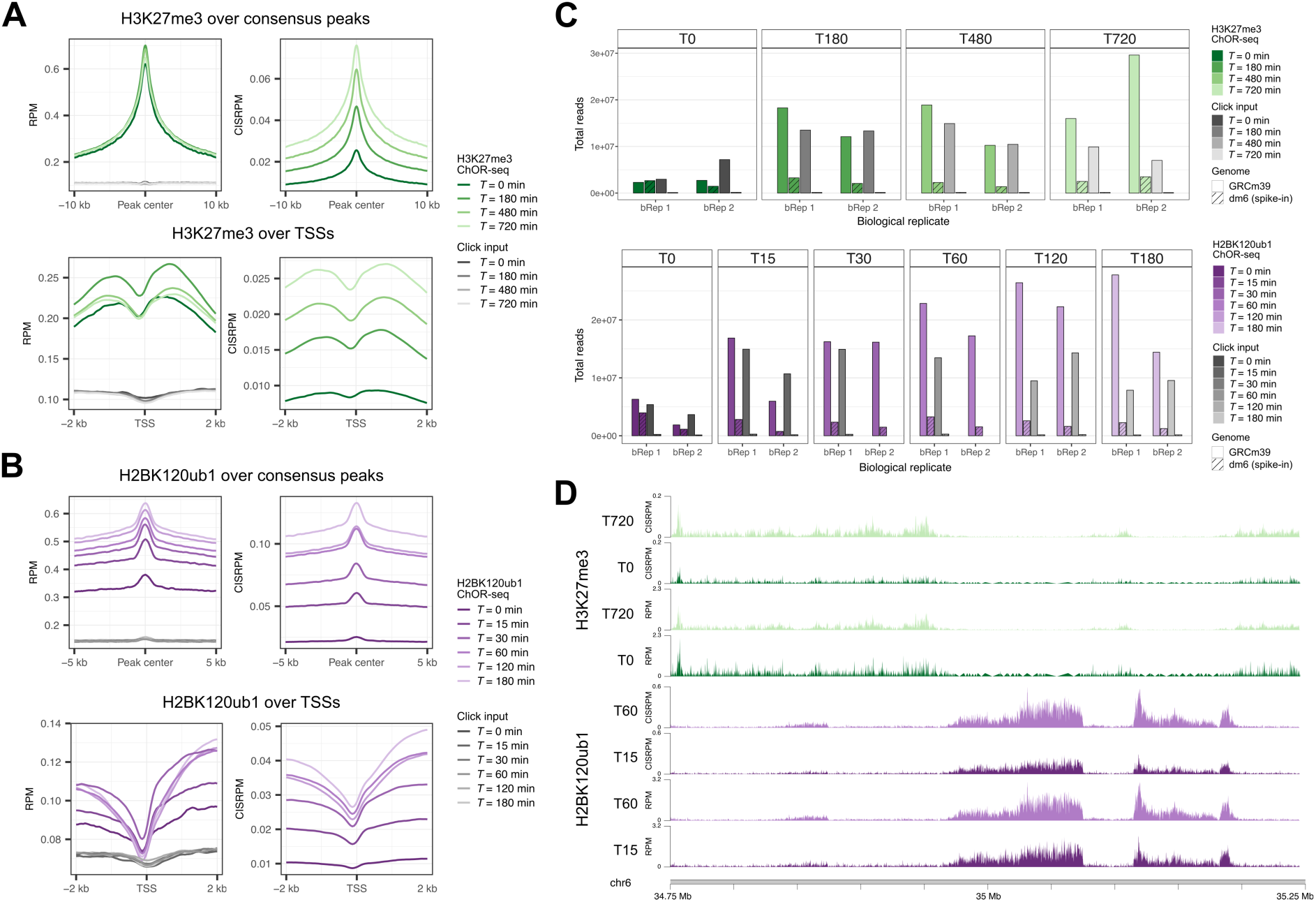
Example of CREPAS’s output for H3K27me3 and H2BK120ub1 ChOR-seq data analysis. **(A)** Profiles of RPM- and CISRPM-normalized H3K27me3 occupancy signal across consensus peak centers (top) and TSSs (bottom). Each line represents the average of biological replicates for that timepoint. **(B)** Profiles of RPM- and CISRPM-normalized H2BK120ub1 occupancy signal across consensus peak centers (top) and TSSs (bottom). Each line represents the average of biological replicates for that timepoint. **(C)** Total mapped and quality-filtered reads for H3K27me3 (top) and H2BK120ub1 (bottom). The bars are grouped by timepoint and subdivided by biological replicate. The exogenous (spike-in) read totals for both the ChOR and EdU input control samples are shown as striped bars, while the endogenous reads are plotted as solid bars. **(D)** Coverage tracks comparing two timepoints of H3K27me3 (top) and H2BK120ub1 (bottom) occupancy after RPM or CISRPM normalization. Each track represents the average of biological replicates for that timepoint.

While RPM normalization adjusts for differences in sequencing depth (total read counts) between samples, it does not allow for quantitative comparisons between samples with variable EdU input control and spike-in total levels (**Figure 4C**). Thus, we used the CISRPM-normalized signal, which is also calculated by CREPAS, to account for the total number of exogenous reads in each sample and any minor variation in spike-in cell mixing in different biological replicates [69]. With this normalization method, the signal across peaks and genes at different time points can be clearly distinguished, whereas with RPM these differences may be underestimated (**Figure 4A-B**). The benefits of this normalization are also evident when contrasting at the genome-wide level, as seen in the coverage tracks of the different timepoints generated by the pipeline (**Figure 4D**). Moreover, the CISRPM-SOI normalization method can correct for when EdU input signal is uneven across the genome, i.e., serves as a bin-by-bin normalization.

The full CREPAS workflow on ChOR-seq data, including preprocessing, alignment, quality control, normalization, and coverage analysis, required less than 22 hours (586 CPU hours) and 1 TB of storage for the H2BK120ub1 dataset, and less than 12 hours (304 CPU hours) and 0.7 TB of storage for the H3K27me3 dataset. Approximately 0.5 TB (H2BK120ub1) or 0.4 TB (H3K27me3) of temporary files could be deleted unless the user intends to resume the workflow from intermediate steps.

### CREPAS facilitates reproducible SCAR-seq and OK-seq analyses to study histone mark segregation to replicated strands

We then tested CREPAS on published SCAR-seq data for H3K27me3 and H3K27ac, generated from the MCM2 histone-binding domain mutant [38]. This mutation prevents the protein from binding parental H3–H4 and causes a strong strand bias, with old histones preferentially segregated to leading strands and new histones to lagging strands [17]. This asymmetric inheritance disrupts the faithful transmission of histone modifications such as H3K27me3 and H3K27ac during mitosis and leads to global epigenomic alterations [38].

Our analysis of the H3K27me3 and H3K27ac SCAR-seq samples with CREPAS reproduced these results, as shown in the plots that summarize the partition signal around initiation zones (**Figure 5A-B**). H3K27me3 showed a strong leading strand bias in nascent chromatin of MCM2-2A cells. This asymmetry decreased over time, although substantial H3K27me3 asymmetry was present 8 h post-replication (**Figure 5A**). To determine the replication initiation zones, we also used CREPAS to reanalyze an OK-seq dataset from mESCs [17]. In contrast to H3K27me3, H3K27ac showed a clear lagging strand bias in MCM2-2A nascent chromatin.

**Figure 5.**
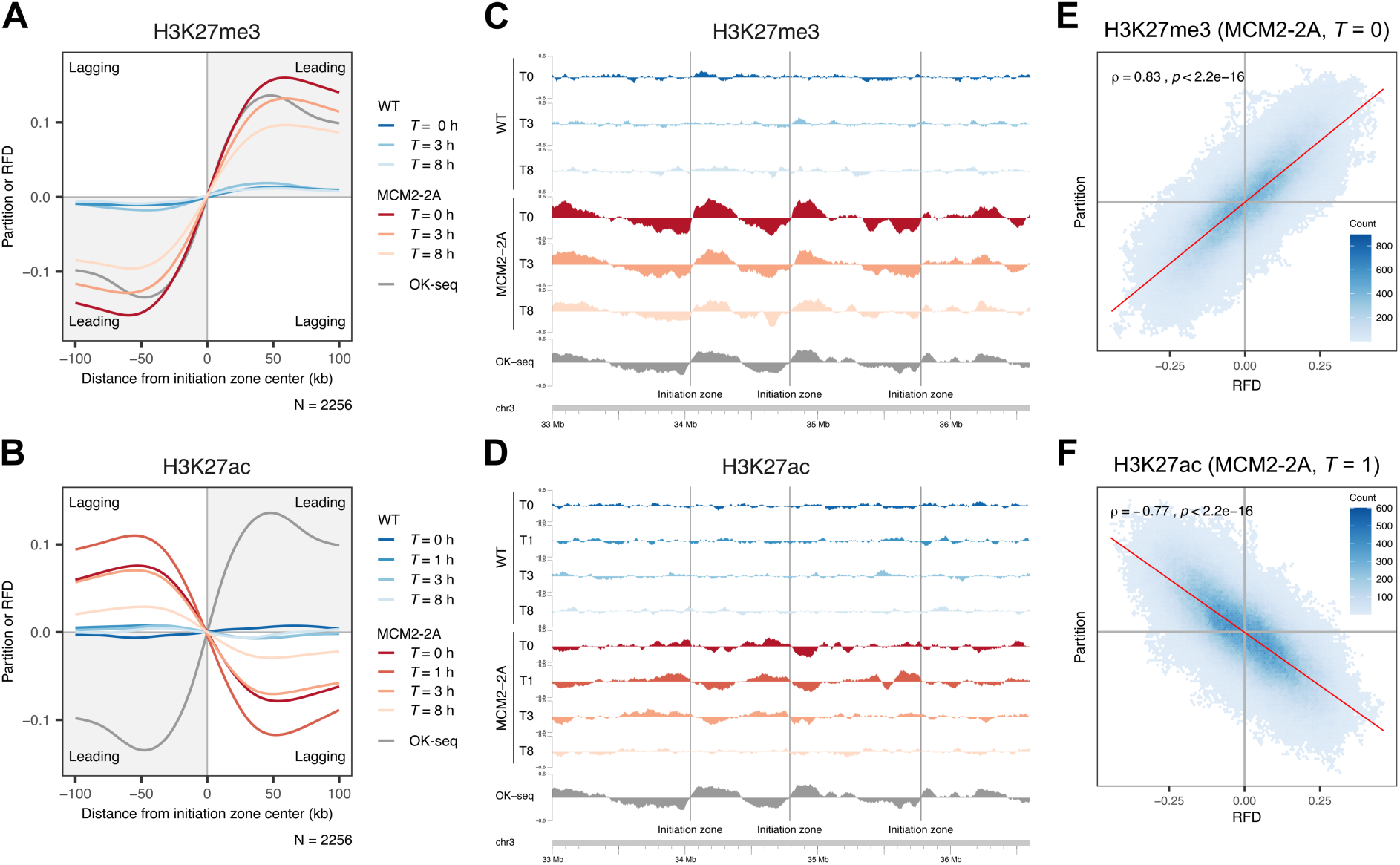
Example of CREPAS’s output from H3K27me3 and H3K27ac SCAR-seq and OK-seq data analysis. **(A and B)** Average smoothed SCAR-seq profile, showing replication fork directionality (RFD, measured by OK-seq), and asymmetry (measured as partition to leading and lagging strand), for H3K27me3 (A) or H3K27ac (B) across replication initiation zones (N). **(C and D)** Representative SCAR-seq profiles showing partitioning of H3K27me3 and H3K27ac between sister chromatids and RFD (OK-seq, grey) in wild-type (WT) and MCM2-2A mutant cells across timepoints. Vertical grey lines denote initiation zones. **(E and F)** Binned scatter plots of H3K27me3 and H3K27ac SCAR-seq partition and RFD show association between histone segregation and RFD. Two-sided Spearman’s rank correlation coefficient and red line representing a linear regression fit to the data are shown.

The asymmetry strengthened during the first hour after replication and then gradually resolved (**Figure 5B**), consistent with a transient post-replicative wave of H3K27ac on new lagging-strand histones [38]. In MCM2-2A cells, this asymmetry persisted even 8 h after replication. Additionally, CREPAS can be used to generate coverage tracks and explore the partition and RFD signal at specific genomic regions of interest (**Figure 5C-D**). Similarly, correlating the partition score with the measurements of RFD by OK-seq also allows for assessment of the old and new histone distribution to the leading and lagging strands (**Figure 5E-F**).

The full CREPAS workflow on SCAR-seq data, including preprocessing, alignment, quality control, normalization, and partition analysis, required less than 6 hours (81 CPU hours) and 0.7 TB of storage for the H3K27me3 dataset, and less than 8 hours (126 CPU hours) and 1.1 TB of storage for the H3K27ac dataset. The OK-seq reprocessing required less than 2 hours (8 CPU hours) and 0.2 TB of storage. Approximately 0.3 TB (H3K27me3), 0.5 TB (H3K27ac) or 0.18 TB (OK-seq) of temporary files could be deleted unless the user intends to resume the workflow from intermediate steps.

## DISCUSSION

Sequencing-based approaches have substantially advanced our understanding of replication-coupled chromatin dynamics, revealing how chromatin accessibility and protein occupancy are restored on nascent DNA. Methods such as repli-ATAC-seq and ChOR-seq have mapped these processes genome-wide, and strand-specific assays like SCAR-seq have added directional resolution, enabling quantitative assessment of chromatin maturation on the leading and lagging strands. As the diversity of nascent chromatin assays increases, the field increasingly depends on computational frameworks capable of integrating heterogeneous datasets, including multiple chromatin-interacting factors and post-translational modifications, while ensuring reproducibility and accessibility for a broad research community. Existing pipelines such as nf-core/chipseq [22], nf-core/atacseq [23], SnakePipes [27] or GenPipes [25] provide valuable foundations for analyzing steady-state chromatin features, but no single workflow adequately addresses the complexity of information emerging from nascent DNA assays. Continued development of modular, scalable, and transparent analysis tools is therefore essential to fully leverage these technologies and advance our understanding of epigenome maintenance across DNA replication.

CREPAS builds on this need by adopting a philosophy of modular, well-defined workflows implemented in Nextflow DSL2, enabling clear separation of analytical steps, straightforward maintenance, and flexible customization by users [92]. CREPAS is portable across compute environments, including cloud platforms and high-performance clusters, and it is fully containerized, ensuring easy installation and identical behavior across systems. Its resource-efficient execution model allows users to process nascent DNA datasets at scale, even when analyzing multiple histone marks or chromatin-associated factors from different assays.

To understand the mechanisms that drive epigenetic memory, including histone recycling and post-replication chromatin maturation, it is essential to fully leverage data from replication-aware assays at locus resolution. CREPAS provides a standardized and efficient framework for analyzing data from quantitative and time-resolved methods such as qChIP-seq, qChOR-seq, and SCAR-seq. By incorporating spike-in normalization and downsampling strategies, CREPAS enables the quantitative profiling of epigenetic factors across multiple time points, capturing changes in protein levels and histone mark abundance as chromatin matures following replication. Integration of these datasets with OK-seq and SCAR-seq further links epigenetic inheritance to replication dynamics and allows the restoration of chromatin features to be resolved at specific genomic loci and on individual daughter strands.

The workflow produces standardized outputs, including normalized signal tracks, strand-specific enrichment profiles, quality metrics, and summary plots and reports that facilitate quantitative comparisons, as well as downstream interpretation and integration with additional genomic datasets. While CREPAS reduces the labor of configuring, maintaining, and reproducing complex analyses, careful parameter selection and biological interpretation remain essential, particularly when working with assays that differ in target, temporal resolution, labeling strategies, or enrichment characteristics.

## ACKNOWLEDGEMENTS

We thank the staff of the CPR/reNEW Genomics Platform for their excellent support and the members of the Groth laboratory for fruitful discussions.

## AUTHOR CONTRIBUTIONS

S.R.P.: Conceptualization, data curation, formal analysis, investigation, methodology, software, visualization, writing – original draft, writing – review and editing. Q.D.: Investigation, writing – review and editing. A.B.: Investigation, writing – review and editing. A.G.: Conceptualization, funding acquisition, project administration, supervision, writing – review and editing. N.A.: Conceptualization, data curation, formal analysis, project administration, supervision, writing – original draft, writing – review and editing.

## DATA AVAILABILITY

CREPAS is available at https://github.com/grothlab/crepas and at https://doi.org/10.5281/zenodo.20788108. Previously published ChOR-seq H3K27me3 and H2BK120ub data are available at GSE204981. H3K27me3 and H3K9me3 ChIP-seq data are available at GSE154379. H3K27me3 and H3K27ac SCAR-seq data are available at GSE154380. OK-seq data are available at GSM3290342.

