## Supplementary Figure S1 for "CREPAS: a reproducible nascent chromatin sequencing analysis pipeline for epigenome replication studies"

### SUPPLEMENTARY DATA

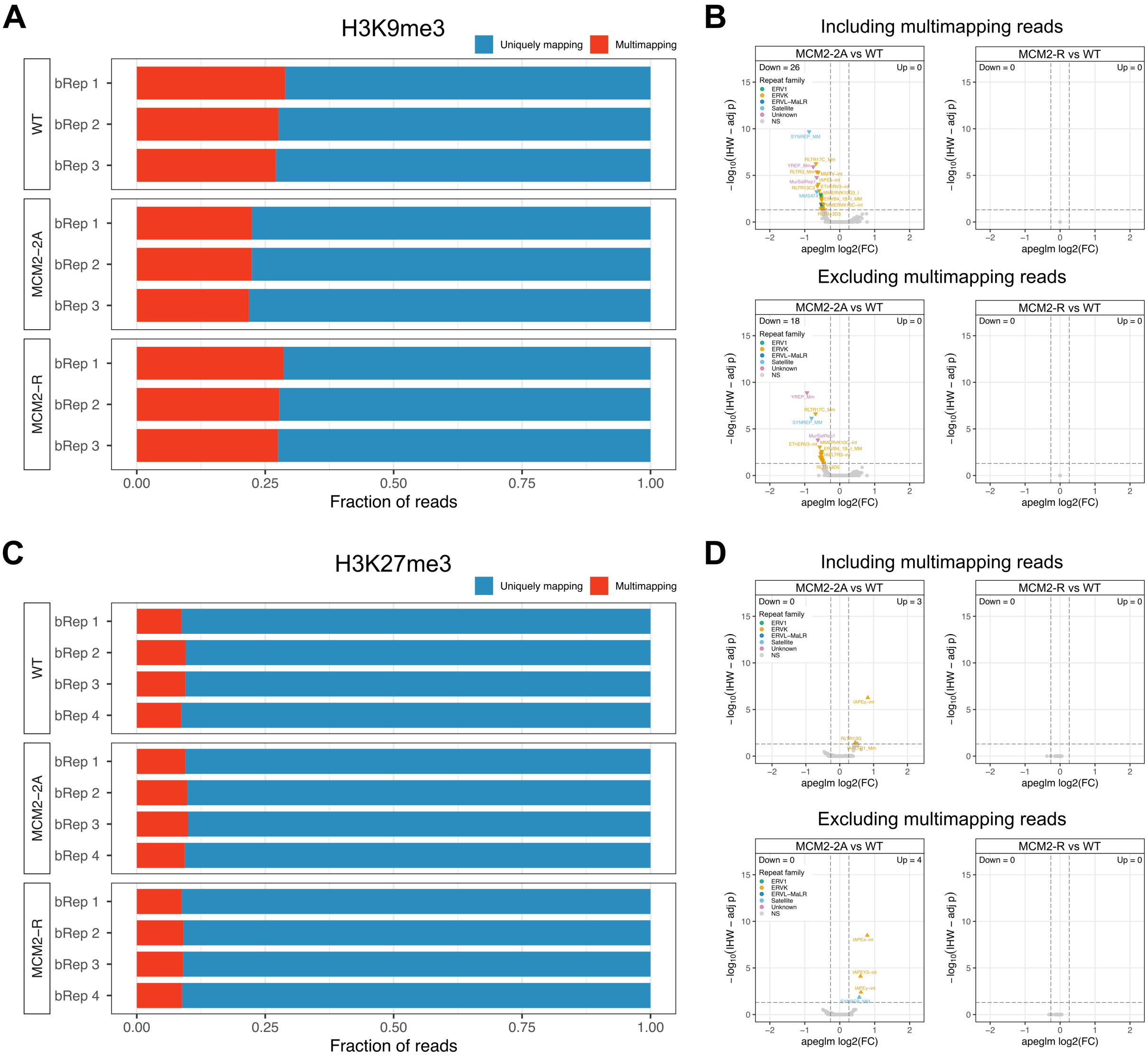


**Supplementary Figure 1. Multimapping read and transposable element quantification in ChIP-seq data with CREPAS facilitates the detection of histone mark occupancy gains and losses at repetitive elements. (A)** Fraction of multimapping reads in H3K9me3 ChIP-seq. **(B)** Repeat subfamilies with significant H3K9me3 loss in MCM2-2A mutant and MCM2-R rescue compared to wild-type (n = 3 biological replicates per genotype, two-tailed threshold-based Wald tests, lfcThreshold = $\log_2(1.2)$, independent filtering and p-value adjusted by independent hypothesis weighting with a significance cutoff $\alpha = 0.05$. **(C)** Fraction of multimapping reads in H3K27me3 ChIP-seq. **(D)** Repeat subfamilies with significant H3K27me3 loss in MCM2-2A mutant and MCM2-R rescue compared to wild-type (n = 4 biological replicates per genotype).
